# DNA methylation is not a driver of gene expression reprogramming in young honey bee workers

**DOI:** 10.1101/2021.03.12.435154

**Authors:** Carlos A. M. Cardoso-Junior, Boris Yagound, Isobel Ronai, Emily J. Remnant, Klaus Hartfelder, Benjamin P. Oldroyd

## Abstract

Intragenic DNA methylation, also called gene body methylation, is an evolutionarily-conserved epigenetic mechanism in animals and plants. In social insects, gene body methylation is thought to contribute to behavioral plasticity, for example between foragers and nurse workers, by modulating gene expression. However, recent studies have suggested that the majority of DNA methylation is sequence-specific, and therefore cannot act as a flexible mediator between environmental cues and gene expression. To address this paradox, we examined whole-genome methylation patterns in the brains and ovaries of young honey bee workers that had been subjected to divergent social contexts: the presence or absence of the queen. Although these social contexts are known to bring about extreme changes in behavioral and reproductive traits through differential gene expression, we found no significant differences between the methylomes of workers from queenright and queenless colonies. In contrast, thousands of regions were differentially methylated between colonies, and these differences were not associated with differential gene expression in a subset of genes examined. Methylation patterns were highly similar between brain and ovary tissues and only differed in nine regions. These results strongly indicate that DNA methylation is not a driver of differential gene expression between tissues or behavioral morphs. Finally, despite the lack of difference in methylation patterns, queen presence affected the expression of all four DNA methyltransferase genes, suggesting that these enzymes have roles beyond DNA methylation. Therefore, the functional role of DNA methylation in social insect genomes remains an open question.

## Introduction

DNA methylation is a reversible chemical modification of DNA whereby methyl groups are added to cytosines in CpG dinucleotides by enzymes of the DNA methyltransferase family (DNMTs). In mammals DNA methylation plays important roles in regulating gene expression, including inactivation of the X chromosome in females, transposon suppression and genomic imprinting, a mechanism by which parents influence gene expression in offspring (1). The role of DNA methylation in genomic regulation of invertebrates is less clear, in part because the two main model species, the fruit fly *Drosophila melanogaster* and the nematode *Caenorhabditis elegans*, either lack methylation (*C. elegans*) or have extremely low and transient levels of methylation (*D. melanogaster*) (2, 3). In contrast, the honey bee (*Apis mellifera*) genome project revealed a functional epigenetic system comparable to that of vertebrates (4, 5). Since this discovery the honey bee has emerged as a model species for epigenetics studies in invertebrates, as, in contrast to mammals, the methylation marks in honey bee genomes are sparse and mostly restricted to gene bodies (intragenic DNA methylation) (2, 6, 7). This sparseness brings a technical advantage to researchers, as it facilitates the study of gene body methylation and its potential roles in the regulation of gene expression without noise from methylated cytosines of other genomic compartments, such as promoters and transposons. Mechanistically, it has been proposed that gene body methylation modulates the affinity of cofactor binding in regulatory DNA methylation-dependent regions in order to regulate RNA polymerase II activity (8–11).

In honey bees and other social insects, gene body methylation has been associated with behavioral and phenotypic plasticity (reviewed in (11, 12)). For example, methylome differences have been found between queens and workers in honey bees (6, 13) and ants (7). Differences in gene body methylation were also associated with division of labor among ant (7, 14) and bee (15) workers. Furthermore, DNA methylation has been associated with several biological processes, including aging, reproduction, aggressiveness, response to social stimuli, memory and learning, and the haplodiploid sex determination system (15–25).

In support of the hypothesis that gene body methylation of honey bees modulates transcription, the RNA interference-mediated knockdown of the *DNA methyltransferase 3* (*Dnmt3*) gene, which codes for the enzyme responsible for *de novo* DNA methylation, affected 14% of the honey bee worker transcriptome (9). RNA splicing was also affected, corroborating previous *in-silico* predictions that associated DNA methylation with alternative splicing (6, 7, 13, 15, 26–29). Even more spectacularly, the knockdown of *Dnmt3* in young female larvae resulted in adults with a queen-like phenotype, mimicking the transcriptional program induced by a royal jelly diet (16).

More recent studies, however, now cast doubt on the role of gene body DNA methylation as a flexible regulator of gene expression in social insects, and an emerging consensus is that DNA methylation is genotype-specific (30–41). For example, a reanalysis of the *Dnmt3* knockdown data of Li-Byarlay et al. (2013) suggested that the original analysis had overestimated the number of regulated genes, and that the *Dnmt3* knockdown had in fact only a minor effect on the honey bee gene body methylation pattern, and hence, on gene expression (37). Furthermore, recent studies have not provided support for the proposed association between differential methylation and alternative splicing in honey bees (38) and other social insects (34, 42–44).

Particularly compelling evidence that DNA methylation tends to be sequence-specific rather than representing a flexible gene regulatory mechanism comes from the clonal raider ant (*Ooceraea biroi*) (31). *O. biroi* reproduces asexually, and therefore allows experiments in a uniform genetic background (45). Libbrecht et al. (2016) showed that DNA methylation is not associated with reproductive and asexual stages in *O. biroi* workers, casting doubt on its role in other ants and honey bees. These authors proposed that the previously reported differential methylation patterns seen between experimental groups may have be an artefact arising from combinations of colony-specific methylation patterns.

In this study, we test the hypothesis that differential methylation is associated with differential gene expression in response to environmental change, versus the alternative hypothesis that differentially methylated regions are a colony and/or individual-specific character that does not vary in response to environmental change. To do this we compared the gene expression and methylation patterns in the brains and ovaries of young honey bee workers as they matured in the presence or absence of their queen. Honey bee queens, through their mandibular gland pheromones, influence the behavioral maturation and reproductive capacity of workers (46–48), and this process involves changes in the expression of hundreds of genes in the brain and ovary (49–52), including the *Dnmt* genes (22, 53). Furthermore, instead of generating data from whole body methylomes, we compared methylation patterns in the tissues (brain and ovary) most likely to respond to the absence/presence of a queen. This allowed us to further examine whether differential methylation is related to tissue function, social context or to genotype.

## Results

### Social context does not affect the brain and ovary methylomes of young honey bee workers

We sequenced and analyzed at single base-pair resolution the complete methylomes of brains and ovaries of honey bee workers reared in queenright or queenless colonies. After removing adaptors and reads of low quality, and aligning the methylomes to the honey bee reference genome (54) we obtained high coverage in all sequenced samples (Supplemental Table S1). The conversion rate of bisulfite treatment was above 99.4% (Supplemental Table S1), indicating a low frequency of false-positive methylated CpGs. The observed frequency of methylated CpGs (∼1% - Supplemental Table S2) is consistent with previous studies on honey bees and other holometabolous insects (6, 7, 15, 55).

With these data we were able to ask whether the methylomes of workers differ when they are in a queenright (QR) or queenless (QL) social context, especially in the two tissues that are known to respond transcriptionally to the presence of the queen: the brain and the ovary (49–51). Comparing three pairs of QR colonies with three pairs of QL colonies we found 879 differently methylated regions (DMRs) for the brains of QL and QR workers (Supplemental Table S3) and 376 DMRs for the ovaries (Supplemental Table S4). However, we found that the number of DMRs shared by two or more colonies was very low for both tissues (Fig. 1A). This indicates that the majority of candidate DMRs previously identified (e.g., Supplemental Table S3 and S4) were driven by between-colony variability and do not reflect a reproductible effect of ‘social context’.

**Figure 1.**
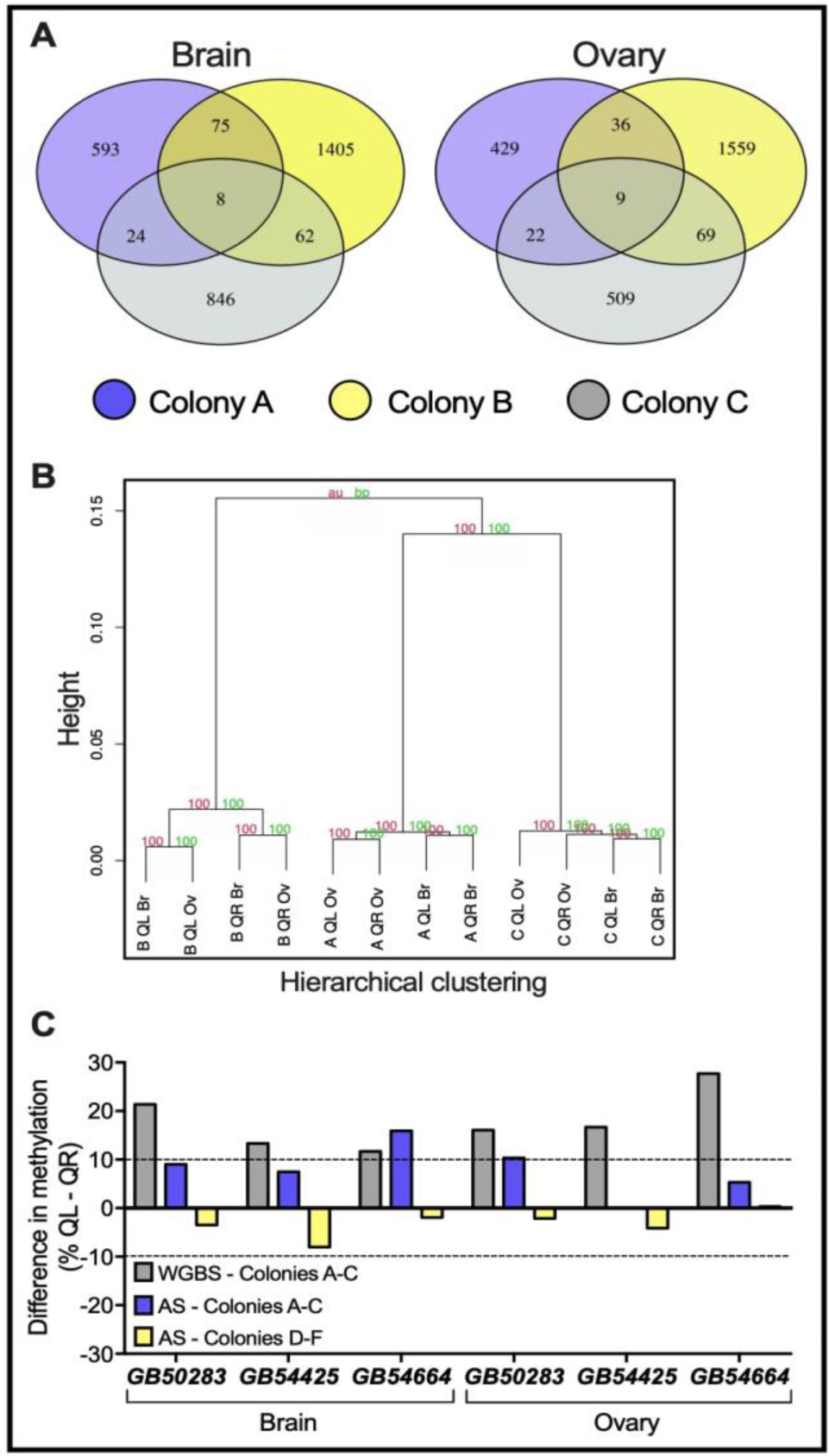
Social context does not drive significant alterations in the methylomes of honey bee workers. (*A*) Number of DMRs comparing QR and QL methylomes across all three colonies assessed for WGBS. Intersections show the number of DMRs shared by more than one colony. Hierarchical clustering showing the correlation-based distances of methylation patterns across all WGBS samples; the numbers show high support values for both approximately unbiased (au) and bootstrap probability (bp) statistics. Samples are identified by their colony origin (A, B or C), social context (QR or QL) and tissue (Br – Brain, Ov – Ovary), respectively. Methylation frequency (%QL - %QR) of three top-ranked DMRs identified in the brain and ovaries for the six colonies. The gray bars represent the frequency of methylated sites between QL and QR bees obtained from whole-genome bisulfite sequencing (WGBS), and the blue bars for the amplicon bisulfite sequencing (AS) for the original colonies A-C. The yellow bars show the frequency of methylated sites (%QL - %QR) obtained from amplicon sequencing of samples from three additional colony pairs (colonies D-F). The dashed lines represent the 10% threshold for considering a region as differentially methylated. Data from individual cytosines and coverage can be found in Supplemental Fig. S2 and Supplemental Table S5, respectively.

Hierarchical clustering of worker methylomes showed that samples from the same colony cluster together, irrespective of social context (Fig. 1B). When comparing the number of DMRs in the methylomes of the three colonies regardless of social context (i.e., A *vs*. B, A *vs*. C, and B *vs*. C), we found over 10,000 significant DMRs in the pairwise colony comparisons (Supplemental Fig. 1). Therefore, the effect of ‘colony’ on the worker methylomes is of much greater magnitude than the effect of ‘social environment’, if there is such an effect.

For a more in-depth analysis, we next selected three top-ranked candidate DMRs for both tissues from the QR *vs*. QL comparison for amplicon sequencing (Fig. 1C, Supplemental Table S5). Bisulfite-treated DNA extracts from the same colonies as those used for WGBS (colonies A-C) were PCR-amplified and sequenced in a high-throughput platform. To increase sample size and check for data reproducibility we also added samples from three new pairs of colonies (colonies D-F) to this analysis. The high-coverage obtained by amplicon sequencing (average >10,000 reads/region, range 672-30,823, Supplemental Table S5) suggests that the majority of cell types present in the brain and ovary tissues are represented in this dataset. Amplicon sequencing revealed that the methylation differences (QR *vs*. QL) observed in the original WGBS data (Fig. 1C – grey bars) did not reach the differential methylation threshold set at 10%, despite the increased coverage for the original samples (Fig. 1C – blue bars, Supplemental Fig. 2; Supplemental Table S5), or samples from the three new independent colonies (Fig. 1C – yellow bars). Importantly, the differential methylation pattern for the majority of regions (five out of six) from colonies D-F was in the opposite direction to that of colonies A-C. Together, these comparisons provide strong support for the hypothesis that the differences seen in the WGBS comparisons of the QR *vs*. QL workers were indeed a result of strong differences in the methylomes of colonies A-C, and not a consistent consequence of social context, either in the original colonies, nor in the three new colonies.

### Differential methylation is not associated with differential gene expression when comparing different colonies and social context

To further investigate whether DNA methylation-mediated transcriptional responses are triggered by queen exposure, we determined the expression patterns of nine genes displaying DMRs found in the contrast of the QR *vs*. QL methylomes (Supplemental Tables S3, S4). We found that only three differentially methylated genes (*GB42836, GB49839* and *GB54664*), were differentially expressed (Fig. 2, Supplemental Table S6), and there was no differential expression for the other six genes in the respective tissue displaying differential methylation. This lack of correlation between differential methylation and differential expression was also statistically confirmed (Supplemental Fig. S3, Pearson correlation r = 0.37, *p* = 0.23, n = 12). Taken together, although major transcriptional responses to the presence or absence of a queen in a colony are regularly observed and reported for the brain and ovary of workers (49–52), our data indicate that there is no direct association between the DNA methylation status of a gene and its differential expression, even in the context of a very dramatic contrast in social context.

Given that the genome methylation differences appeared to be much more related to colony genotype than to social context (Fig. 1 and Supplemental Fig. 1), we next asked whether colony-specific methylation patterns are associated with differential gene expression between colonies. To do so, we first estimated the “colony effect” by ascertaining how many regions showed different methylation levels in the brain and ovary of workers, regardless of the social context (Supplemental Fig. S1). We found large numbers of DMRs in both the brain (n = 3,603) and ovary (n = 1,997) that differed in their methylation level by at least 10% when comparing the same genomic region pairwise among the colonies (Fig. 3A). This indicates that there is a core set of genomic regions that are hypervariable with respect to their methylation status. Interestingly, these regions are not chromosome specific (Supplemental Fig. S4), suggesting a colony-specific methylation fingerprint across all chromosomes. Also, almost 60% of the hypervariable regions (n = 1,194) seen in the ovaries were the same as those found in the brains (Fig. 3B), suggesting that these colony-specific signatures are independent of tissue.

**Figure 2.**
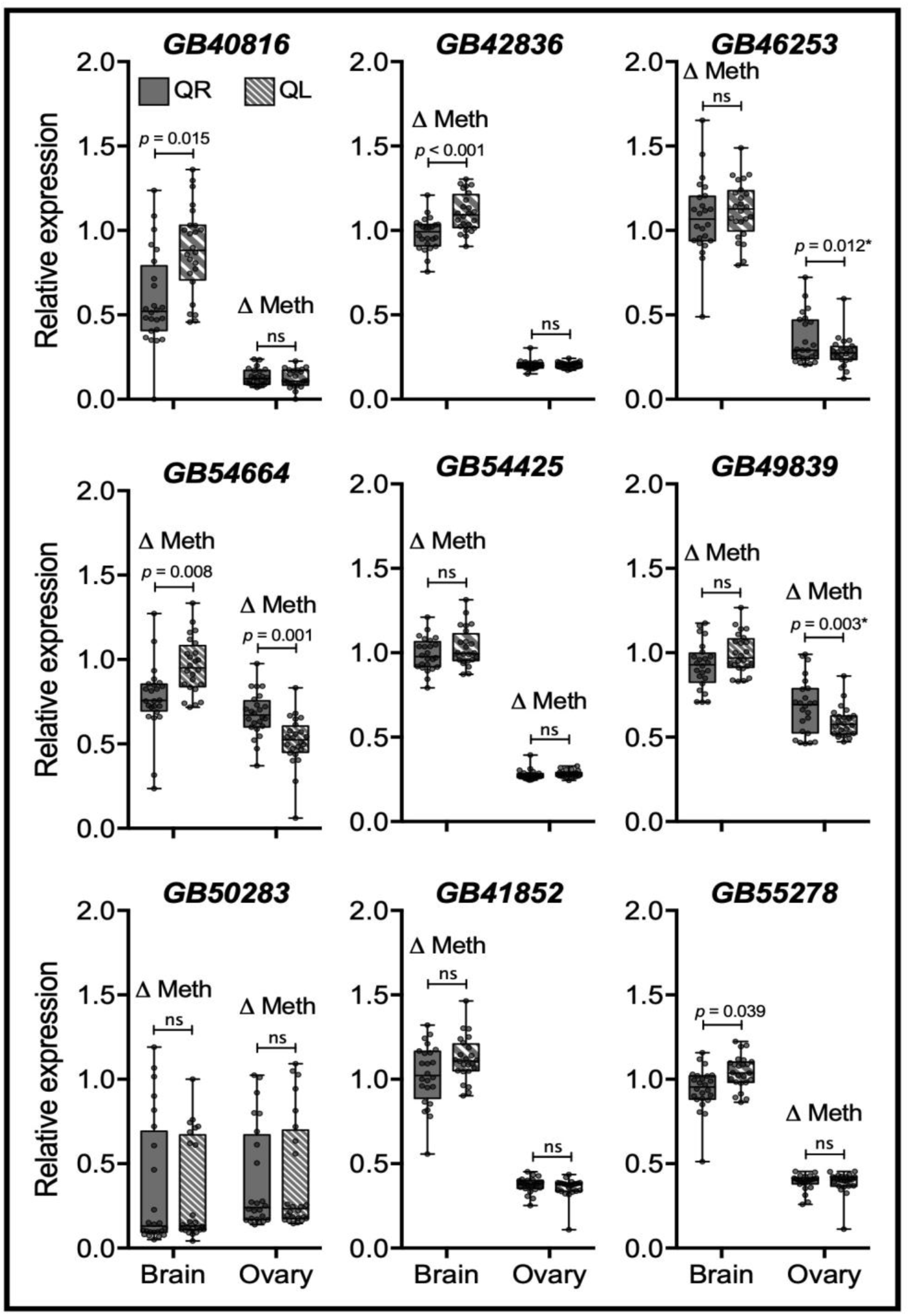
Relative gene expression analysis of nine differentially methylated genes in the brain and ovaries of workers from queenright (QR) and queenless (QL) colonies. “Δ Meth” shows the tissue where DNA methylation was affected by social context (Supplemental Tables S3, S4). Note that only *GB42836* (brain), *GB49839* (ovary) and *GB54664* (both tissues) were both differently methylated and expressed in the QR *vs*. QL contrast. Each box shows the interquartile range (25^th^-75^th^ percentiles) and the median (line), while whiskers represent the minimum and maximum values. Gray dots inside boxes represent individual samples (n = 3 colonies, 8 samples per colony, 24 in total). Statistical information: GLMM test of differences between means with Tukey correction for multiple pairwise comparisons, an asterisk (*) represents the genes with significant difference in the post-hoc test but non-significant for the main ‘social context’ effect, ‘ns’ indicates *p* > 0.05, see Supplemental Table S6 for further information.

**Figure 3.**
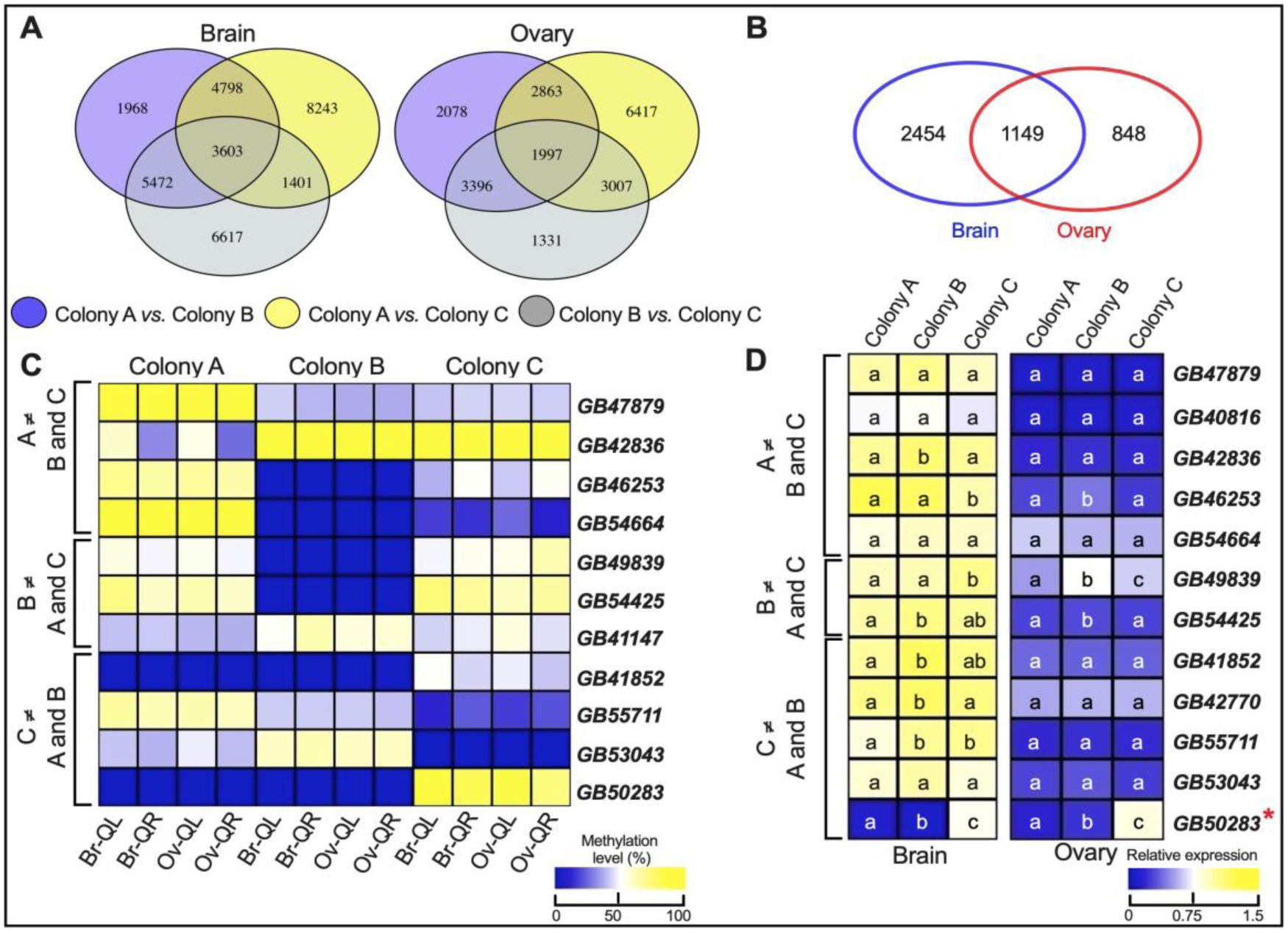
Differences in the methylomes and gene expression levels between source colonies. (*A*) Venn diagrams for the number of DMRs in the brain and ovaries comparing three different colonies. Numbers inside the intersections represent the regions that show differential methylation of at least 10% in one source colony compared to the other two source colonies. For example, the intersect between blue and yellow region (e.g., 4798 DMRs for the brain or 2863 DMRs for the ovaries) represents the regions from colony A that differ in methylation level by at least 10% from the same regions in colony B and C. The intersection of all three comparisons, i.e. the central intersect, represents the regions with three degrees of DNA methylation levels (for example, 0% methylation level for colony A, 20% for colony B and 70% for colony C). (*B*) Number of DMRs from central intersects displayed in Fig. 3A shared by the brain and ovary tissues. (*C*) Heatmap showing colony-specific methylation validated by high-throughput amplicon sequencing of 11 DMRs. For this analysis, we used the WGBS data to select genes that had a specific pattern for one colony (hypermethylated or hypomethylated), but differed from the patterns observed in the other two colonies. Methylation level for colonies A-C are shown as follows: Br-QL – brain queenless; Br-QR – brain queenright; Ov-QL – ovary queenless; Ov-QR – ovary queenright. Coverage of each region can be found in Supplemental Table S5 and methylation level of individual CpG sites are displayed in Supplemental Figs. S2, S6, S7. (*D*) Colony-specific gene expression of differentially methylated genes. Note that only one gene, *GB50283* (red asterisk), had a methylation pattern (Fig. 3C) that seemed correlated with its gene expression level. Note that the list of genes shown in Fig. 3D is the same as in Fig. 3C, the exceptions being the gene *GB41147*, a differentially methylated gene which relative expression was not examined, and the genes *GB40816* and *GB42770*, which were differently methylated in the WGBS data but not validated by amplicon sequencing. Different letters inside the heatmap boxes indicate statistical differences between colonies (GLMM test, n = 16 samples per colony and tissue). Differential expression between tissues was observed for all of the tested genes (Supplemental Table S7), with the exception of *GB54664*.

Next, we asked whether changes in DNA methylation were associated with differential gene expression between colonies. For these analyses we selected 11 genes that were either hypermethylated or hypomethylated in one source colony, but methylated in the opposite direction in the other two source colonies (e.g., hypermethylated in colony A but hypomethylated in colonies B and C). We confirmed the methylation pattern of these regions by amplicon sequencing of colonies A-C (Fig. 3C, Supplemental Table S5) and D-F (Supplemental Fig. S5, Supplemental Table S5). We noticed that the methylation patterns of a given genomic window were reasonably stable across different social contexts and tissues (Supplemental Figs. S2, S6). These analyses revealed strong colony-specific methylation patterns for all six analyzed colonies. Thus, if DNA methylation plays a role in regulating gene expression, these genes would be strong candidates for differential gene expression, as their differential methylation across colonies was much greater than the differences we observed for social context.

Only one gene (*GB50283*) out of the 12 genes assessed, which was strongly methylated in colony C compared to the other two colonies (Fig. 3C), turned out to be differently expressed among colonies (Fig. 3D, *p* < 0.01, Supplemental Table S7). For the other genes assessed, we found that even clear differences in DNA methylation profiles between colonies (Fig. 3C) did not affect their expression (Fig. 3D). After all these comparisons between different colonies and radically different social contexts, we conclude that there is little evidence in support of the hypothesis that differences in DNA methylation drive alterations in gene expression in the two tissues that are most likely to respond to social context. Hence, the brain and ovary methylomes of young honey bee workers are primarily a manifestation of colony identity rather than a mediator between social environmental changes and gene expression.

### Brain and ovary methylomes are highly similar, despite the functional differences between the two tissues

After showing that DNA methylation of young honey bee workers is colony specific rather than influenced by the social context, we investigated whether there may be tissue-specific differences between the brain and ovary methylomes. Comparing the methylomes of tissues with completely distinct biological functions, we expected to identify hundreds of DMRs associated with the transcriptomic differences between these two tissues. Surprisingly, we found only nine DMRs across all three source colonies when comparing the methylomes of the two tissues, irrespective of social context (Fig. 4A). These regions are associated with only four genes: *GB47277, GB51802, GB55278*, and *GB50784*. The amplicon sequencing analysis performed for two of these DMRs confirmed the differences originally seen in the WGBS data for these two tissues (Fig. 4B and Supplemental Fig. S7).

**Figure 4.**
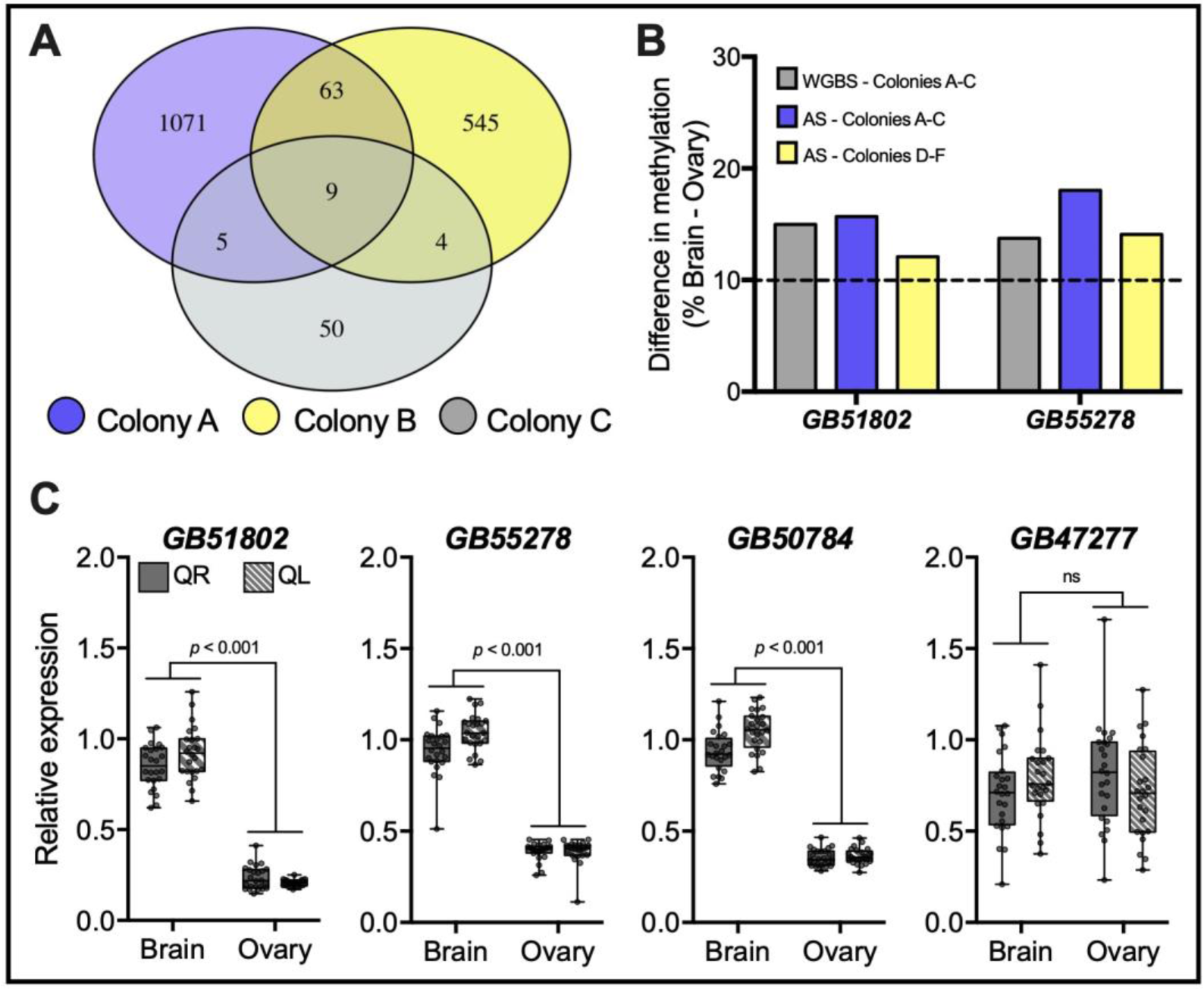
Differences in the methylomes and gene expression levels between brain and ovary tissue. (*A*) Venn diagram shows the number of DMRs in the contrast between brain vs. ovary methylomes in all three colonies (A-C). These nine DMRs are associated with four genes. (*B*) Difference in methylation for two of the four genes seen in the WGBS data (gray bars) and validation by amplicon sequencing in colonies A-C (blue bars) and D-F (yellow bars). The dashed line represents the 10% threshold for considering a region as differentially methylated. Coverage of each region can be found in Supplemental Table S5 and methylation level of individual CpG sites in Supplemental Fig. S7. (*C*) Transcript levels of the four differentially methylated genes quantified by quantitative PCR in the brain and ovary of young honey bee workers. Note that *GB55278* was also found to be differentially methylated between the ovaries of QR and QL workers (Fig. 2 and Supplemental Table 4). Each box shows the interquartile range (25^th^-75^th^ percentiles) and the median (line), while whiskers represent the minimum and maximum values. Sample size and statistical analysis are the same as in Fig. 2 and “ns” indicates *p* > 0.05.

When testing whether alterations in gene expression between the brain and ovaries were associated with differences in DNA methylation levels we found that three of the four differentially methylated genes were also differentially expressed (Fig. 4C, Supplemental Table S6). However, differential expression between tissues were identified for 11 out of the 12 genes previously analyzed for colony specificity (Fig. 3D, Supplemental Table S7). Thus, it is not yet clear whether the alterations in methylation level promotes causative alterations in gene expression between the two tissues. From an overall perspective the methylomes of brains and ovaries were strikingly similar, with significant differential methylation seen for only four genes.

### Expression of the four *Dnmt* genes is influenced by social context

As our data strongly indicate that the presence or absence of a queen does not drive significant alterations in the gene body methylation patterns in the brain and ovaries of young honey bee workers (Fig. 1) we wondered what the expression patterns of the genes encoding the four honey bee DNA methyltransferases would look like, especially since previous studies have used the expression of *Dnmt* genes as an indicator of epigenetic events in social insects (16, 18, 20, 23, 25, 56). Furthermore, we have recently reported that the queen mandibular pheromone upregulated the expression of three *Dnmt* genes (*Dnmt1b, Dnmt2* and *Dnmt3*) in the brain of caged honey bee workers (53). Here we report that the expression of all four *Dnmt* genes predicted in the *A. mellifera* genome is affected by the presence or absence of a queen in the colony (Fig. 5, Supplemental Table S6), either in the brain or the ovary. Interestingly, the same social cue (presence/absence of a queen) that upregulated two *Dnmt* genes (*Dnmt1b* and *Dnmt3*) in the brain of young workers had an opposite effect on the expression of three *Dnmt* genes (*Dnmt1a, Dnmt2* and *Dnmt3*) in the ovary. This suggests that expression of the *Dnmt* genes responds to social cues in both tissues, even though their differential expression apparently is not associated with alterations in the methylomes of honey bee workers.

**Figure 5.**
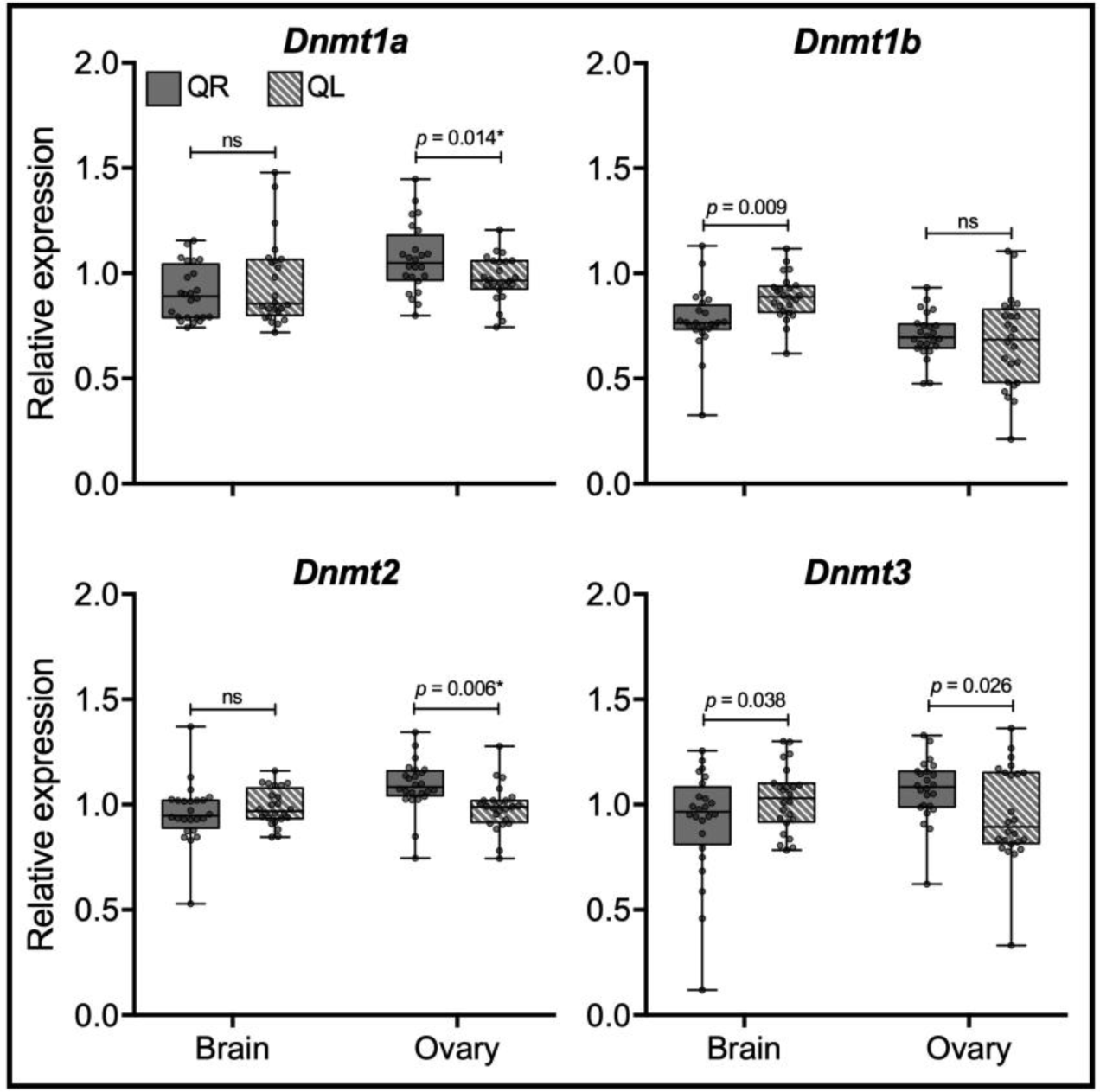
Relative expression of DNMTs encoding genes in honey bee workers kept in two different social conditions, queen presence (QR) or queen absence (QL). Transcript levels were assessed by quantitative PCR in the brain and ovary of young honey bee workers. Each box shows the interquartile range (25^th^-75^th^ percentiles) and the median (line), while whiskers represent the minimum and maximum values. Sample size and statistical analysis are the same as in Fig. 2; “*” represents the genes with a significant difference in the post-hoc test but non-significant for the main ‘social context’ effect, “ns” indicates *p* > 0.05.

## Discussion

With the presence or absence of a queen being one of the strongest contrasts in the social context of a colony, and with DNA methylation as a strong candidate for mediating adaptive responses in the individual colony members, we had expected to see signatures of this social context condition in the methylomes of young honey bee workers. However, all the evidence we obtained suggests the opposite: that gene body methylation is not a major regulator of gene expression reprogramming in young honey bees. First, the contrast between the presence *vs*. absence of a queen was not reflected in significant alterations in the respective methylomes, indicating that differential DNA methylation is not a mediator for the transcriptional consequences of queen presence. Second, despite strong differences in gene expression between brain and ovarian tissues (57), we found that their methylomes are very similar. Third, thousands of DMRs were identified when comparing individuals from different colonies. However, even when differences in methylation were greater than 50%, there was no association between DNA methylation and gene expression, except for one gene (*GB50283*, Fig. 3D). Therefore, these data provide strong support for the hypothesis that intragenic CpG methylation is not a driver of gene expression reprograming in the brain and ovary of young honey bees. This result is actually in line with and likely generalizable for other social insects (31, 32, 34, 41, 43, 44, 58).

In contrast to our findings, two previous studies have suggested that honey bee workers kept under different social conditions exhibit a low number of DMRs, and that these are associated with differential gene expression (15, 17). A plausible explanation (32, 39) for these contrasting results is that previously-observed differences in the methylomes of honey bee workers in response to divergent social stimuli arose as an artifact of genotype-associated methylation variants (33). Indeed, when the colony genetic background was standardized, no differences were found between the methylomes of newly-emerged queens and newly-emerged workers (15). Furthermore, ultra-deep analyses on the dynamics of brain DNA methylation of *Dynactin, Nadrin* and *PKCbp1* genes, indicated that their methylation content are not altered over time, and thereby not implicated in honey bee workers’ aging and behavioral maturation (59). Thus, we conclude that the methylomes of honey bee workers do not respond dynamically to changes in the social milieu (e.g., loss of the queen), whereas gene expression levels do so (49–53).

Our results also show that DNMT-encoding genes are differentially expressed, and thus affected by the presence or absence of a queen. This finding is consistent with previous honey bee studies that showed an effect of queen mandibular pheromones on the expression of *Dnmt* genes (22, 53). However, these alterations seen in the expression of the *Dnmt* genes apparently does not promote a “real-time” epigenetic reprogramming process (this study; Harris et al. 2019), indicating that the expression levels of *Dnmt* genes should not be used as a proxy of possible differences in DNA methylation patterns or levels. This result further highlights the weak association between the expression of *Dnmt* genes, DNA methylation, and gene expression reprogramming seen in honey bees and other insects. For instance, even though the genome of the red flour beetle *Tribolium castaneum* is not methylated (60, 61), the knockdown of a *Dnmt1* ortholog resulted in offspring showing developmental arrest and high mortality (61). Similar developmental defects in response to a knockdown of *Dnmt1* were observed in the reproductive system of females of the milkweed bug *Oncopeltus fasciatus* (37, 62). These phenotypic alterations are not a consequence of transcriptomic changes, even though the *Dnmt1* knockdown successfully reduced the global levels of DNA methylation (37). Combined, these results imply that DNMTs have other, not yet understood functions, that go beyond DNA methylation and its maintenance.

After an unequivocal demonstration that colony-specific methylation patterns are major signatures in the methylomes of honey bee workers (Fig. 3), an interesting question emerges: why do individuals from different colonies exhibit such contrasting differences between their methylomes? While with our current data we are not yet able to fully solve this puzzle, we propose that colony-specific methylation patterns are both functional and genotype-associated; with a minimum effect on transcription (37, 38). Our results show that colony-specific methylation patters are predominantly chromosome and tissue independent (Fig. 3B, Fig. 4. and Fig. S4). Therefore, the functional role of intragenic methylation in honey bees is likely to be associated with basal processes necessary for cell viability, e.g., cell division, genome integrity, and/or cell cycle progression (37, 61–63), rather than representing as a marker of tissue identity (Fig. 4), or acting as a dynamic sensor of changes in the social environment (Fig. 1). The finding that the knockdown of the *Dnmt3* gene function, a key orchestrator of honey bee development, is lethal at early development (16) suggests that DNA methylation is essential to honey bee viability. Furthermore, the colony-specific methylation patterns are, at least in part, sequence-specific (32, 39, 40). Ultra-deep amplicon sequencing analyses identified that a honey bee brain displays only a small repertoire of methylation patterns. This may reflect the small number of maternal and paternal epialleles predicted for polyandrous insects (59). Interestingly, the colony-specific methylation patterns are inherited from drones to their worker daughters (40), suggesting a role for the paternally inherited epialleles to the composition of colony-specific worker methylomes.

In conclusion, all evidence generated in this study goes against the hypothesis that gene body methylation is a driver of gene expression re-programming in adult honey bee workers. With our experimental design, using several colonies of standardized within-colony genetic backgrounds and high-throughput genomic sequencing approaches we had originally expected to provide evidence in favor of this hypothesis and to see defined differentially-methylated gene sets between queen states, with correlated expression patterns. As it has turned out, the conclusion to be drawn is that gene body methylation patterns are essentially colony specific and unaffected by even radical changes in social context. Nonetheless, it is possible that intragenic DNA methylation plays a role in gene expression during other stages of a worker’s life cycle. If so, this is likely be restricted to a small set of genes under specific contexts (16, 35, 37, 38), including the *GB51802, GB55278* and *GB50784* genes that show both tissue-specific methylation patterns and differential expression (Fig. 4C). Our results make it clear that, whatever may be the specific function of gene body DNA methylation in the honey bee, it is now undeniable that there is a strong genotype effect shaping the workers’ methylomes, and that future studies need to consider the possible effect of genotype variants (colony and patriline genotypes) in their experimental design. In this sense, we believe that our work makes a significant contribution to the understanding of the meaning of the enigmatic, but evolutionarily-conserved intragenic DNA methylation in an important invertebrate epigenetic model species, the honey bee. As it stands now, we must say that the functional role of gene body methylation in social insects is still an unsolved issue.

## Material and methods

### Bees and manipulation of the social environment

We used workers from six source colonies. These colonies were headed by queens of standard commercial stock (mostly *A. m. ligustica*) each of which had been instrumentally inseminated with sperm from a single male (64). Hence, the workers within each colony pairs were all full sisters, with a genetic relatedness of 0.75, thus minimizing genotypic heterogeneity within colonies.

Queenright and queenless colonies were prepared by splitting each of the six host colonies into two (n = 12), thus generating colony pairs of the same genetic background but differing completely in terms of social context. Brood frames were equalized among the colony pairs to ensure a similar condition in terms of brood presence, despite the presence or absence of the queen. After splitting the host colonies into queenless and queenright halves at a remote apiary, we moved the splits to the apiary at the University of Sydney, thus reducing the tendency of the queenless workers moving back to the respective unit with the queen. The experiments were conducted during the southern hemisphere summers of 2017 (source colonies A-C) and 2018 (source colonies D-F).

Newly-emerged workers from source colonies A-F were obtained by placing sealed brood frames overnight in separate boxes, in an incubator at 34 °C on the day of colony splitting. The newly-emerged workers were divided into two groups (n = 200 bees per group), paint marked with different colors, and then introduced into their respective pair of queenright and queenless host colonies. After four days, the marked workers were collected, snap frozen on dry ice, and stored at −80 °C. At this age workers are known to respond transcriptionally to the presence or absence of a queen (49, 53). As expected, the queen was seen in all QR host colonies at the time of sampling, but never in the QL host colonies. Samples from six of these colonies (colonies queenright A-C and queenless A-C) were initially used for methylome sequencing and subsequently also for the region-specific analyses, whereas the other six colonies (colonies queenright D-F and queenless D-F) were used for the region-specific analyses only, which served to validate and confirm the reproducibility of the methylome results. The brains and ovaries of the workers were dissected as described elsewhere (53, 65).

### Whole Genome Bisulfite Sequencing (WGBS) and Amplicon Sequencing

We first performed Whole Genome Bisulfite Sequencing (WGBS) to identify which genes were differentially methylated in the samples from the first three queenright and queenless colonies (colonies A-C). For each sample we pooled either eight brains or 20 pairs of non-activated ovaries. Pooling was necessary to obtain a sufficient DNA yield for high-throughput sequencing. This resulted in a total of 12 samples, represented by six brain and six ovary samples for each of the two social contexts (three queenright and three queenless). DNA was extracted with the DNeasy Blood and Tissue Kit™ (Qiagen) and quantified with a Qubit 2.0 Fluorometer system (Invitrogen). Before sequencing, 0.01% (w/w) of unmethylated Lambda DNA (Promega) was added to each sample to be later used for calculating the bisulfite conversation efficiency. For WGBS, the DNA samples were sent to the Beijing Genomics Institute (China), where library construction, bisulfite treatment, and sequencing were performed. Bisulfite treatment was done with the EZ-DNA methylation kit (Zymo Research), and paired-end WGBS was performed on an Illumina NovaSeq platform. Each library was sequenced twice in two separate lanes for high coverage. In the bioinformatics analysis, the data from the two lanes were then merged, as we did not detect relevant differences between the two runs. Data on coverage for each sample are given in Supplemental Table S1.

After processing the WGBS data and identification of DMRs, we performed a paired-end ultra-deep amplicon sequencing analysis (59, 66) for 13 of the DMRs revealed by WGBS, using a total of six colony pairs, the original three pairs (source colonies A-C) plus three new colony pairs (source colonies D-F). DNA extractions, bisulfite conversion, and Lambda DNA spiking was performed as described above. Columns were eluted with 20 μL of ultrapure water, and 1 μL of the eluted solution was used as template in the PCR assays. Bisulfite PCR primers were designed to amplify fragments between 140-300 bp of the forward strand of each differentially methylated region and the control Lambda spike (Supplemental Table S8). Bisulfite-treated DNA was amplified using the KAPA HiFi Uracil^+^ Kit™ (Roche). PCR assays were set up with 5 μL of Kapa Master Mix, 0.3 μL of each primer (forward and reverse – 3 pmol/reaction), 1 μL of bisulfite-treated DNA and 3.4 μL of water for a total reaction volume of 10 μL, and performed with the annealing temperatures of the respective primers (Supplemental Table S8). For multiplexing of the samples into two libraries, Nextera barcodes were added to the 5’ ends of all primers. Amplicons from different primers were pooled to generate a separate library for each sample (24 samples in total: queenright and queenless brains and ovaries from each of the source colonies A-F). Samples were purified, and Nextera paired-end libraries were constructed at the Australian Genome Research Facility (AGRF). Libraries were constructed in duplicate for each sample and sequenced (150 bp paired end) in a single flow cell of an Illumina MiSeq platform. Data from the two libraries were merged for downstream analyses.

### Differential methylation analyses from WGBS data

Quality of the raw data was assessed by FastQC 0.11.8 (http://www.bioinformatics.babraham.ac.uk/projects/fastqc) and reads with quality scores < 20 were removed. Trimming was performed with TrimGalore 0.5.0 (www.bioinformatics.babraham.ac.uk/projects/trim_galore/) with a stringency error of 2 bp. Overall, we removed ∼ 0.5% of the total sequencing reads after checking read quality and trimming of the adaptors (Supplemental Table S1). The remaining reads were mapped onto the honey bee reference genome assembly Amel_4.5 (54) using Bismark 0.16.1 (67) and Bowtie 2 2.3.5.1v (68). The honey bee genome assembly Amel_4.5 was the newest honey bee genome version at the time the bioinformatic analyses were performed. Coverage varied between 27- 46 times across samples (Supplemental Table S1). Methylation calling was performed with Bismark software. We used 10-times coverage as a threshold for adequate sequencing coverage of each cytosine as in Herb et al. (2012), to make our data sets comparable. Methylation levels were assessed by the C/T ratio of converted cytosines to unconverted bases (69). Significantly methylated sites were identified using a binomial probability model that takes into account the bisulfite conversion rate for each sample (Supplemental Table S1) as the probability of success, followed by Bonferroni corrections at the 1% significance level using BWASP (34). We removed methylated CpGs with > 500x coverage to avoid PCR-based bias in the analyses.

MethylKit (70) was used for differential methylation analysis. First, the honey bee genome was partitioned into 200 bp sliding windows (step size = 100 bp). Only windows containing at least four sufficiently covered CpGs (two in each strand) were analyzed. A difference threshold of 10% in the methylation level between pairwise comparisons was applied (Herb et al. 2012). A threshold >10 methylated cytosines (sum for all CpGs inside given window) in at least one of the libraries was used to reduce methylome complexity. The list of DMRs was then FDR-corrected and a q-value <0.01 was considered significant. A gene containing at least one DMR was defined as a differentially methylated gene. Gene annotation was performed with Homer 4.9.1 software (71). Analyses were performed in the *R* environment (R Core Team 2018). Hierarchical clustering distances between methylome samples were determined with the R package “pvclust” (73).

### Differential methylation analysis of amplicon sequencing data

Reads were checked for quality using FastQC 0.11.8 (www.bioinformatics.babraham.ac.uk/projects/fastqc), followed by trimming of adaptors and removal of low quality reads (Phred score < 20) using Trimmomatic (74). Between 85-90% of the reads were retained for each library. Bissulfite-converted DNA sequences of the 13 amplicon regions of interest and the Lambda control sequence were used as templates to generate a Bowtie2 index prior to alignment. Data from each of the two duplicate libraries for each sample were aligned with Bowtie2 2.3.5.1v (68) using paired-end default parameters and then converted to BAM files with Samtools (75). BAM files were imported into Geneious software 10.2.4 (76), and alignments for each amplicon were manually checked for each sample. C-to-T variant frequencies were calculated using the ‘Find variant’ function for all CG sites, with a minimum coverage of 50 and a minimum variant frequency of 2%. Additional SNP variants that were still visible after bisulfite conversion, such as G-to-A polymorphisms, were also recorded. The overall cytosine methylation frequency was determined for each of the four treatment groups (QR brain, QL brain queenless, QR ovary, QL ovary) for all of the 13 amplicons by dividing the total amount of C (methylated cytosines) per the total amount of C+T (total amount of methylated and unmethylated cytosines). The results are presented as percentages, and when appropriate, regions were compared in relation to their colony of origin, social context, and tissue type.

### Gene expression analysis

Each sample consisted of four pairs of non-activated ovaries or one brain (n = 8 per source colony and social context combination). Ovaries needed to be pooled to obtain a sufficient amount of RNA. Brains and ovaries were macerated in TRIzol (Invitrogen). Total RNA was extracted using the Direct-zol™ RNA™ Miniprep kit (Zymo Research) according to the manufacturer’s instructions. Samples were treated with Turbo DNase™ (Thermo-Fisher Scientific), and RNA concentrations were determined using a Qubit 2.0 Fluorometer system (Invitrogen). RNA samples were diluted with ultrapure water to a final concentration of 40 ng/μL (brain) and 15 ng/μL (ovary). We used 142.5 ng of ovary RNA and 600 ng of brain RNA to synthesize cDNA using the SuperScript™ III Reverse Transcriptase Kit (Invitrogen) with Oligo(dT) primer (Invitrogen). Ovary cDNA was diluted to 2 ng/μL due to its lower concentration, while brain cDNA was diluted to 5 ng/μL in ultrapure water.

Relative expression was determined by RT-qPCR assays for total of 20 genes, 12 being hypermethylated or hypomethylated in one source colony, four genes whose methylation patterns differ between tissues and four *Dnmt* genes. Assays were set up with 2.5 μL SsoAdvanced™ Universal SYBR^®^ Green Supermix (Bio Rad), 1.25 pmol of each primer, 1 μL diluted cDNA in a total volume of 5 μL using a CFX384 Real-Time System (Bio-Rad). For each sample we conducted three technical replicates and used their mean as the data point. Cycle conditions were as follows: 95 °C for 10 min followed by 40 cycles of 95 °C for 10 s, annealing temperature (Supplemental Table 8) for 10 s and 72 °C for 15 s. At the end of the PCR cycles a melting curve analysis was run to confirm a single amplification peak. Relative gene expression analyses were performed using the a formula that accounts for primer’s efficiency: E^CqMin-CqSample^ − where “E” (Table S3) is the efficiency of primers, “CqMin” is the lowest Cq value for a given gene and “CqSample” is the Cq of that sample (65, 77, 78). Then, two reference genes (*Rp49* [also known as *Rpl32*] and *Ef1α*) were used to normalize the expression levels of the target genes. These control genes have been previously validated for honey bee quantitative PCR analyses (79) and were stable in our analysis according to the BestKeeper software (80). Primer efficiencies (Supplemental Table 8) were calculated based on an amplification curve of 10 points obtained through serial dilution of mixed cDNA samples. The list of primers used is given in Supplemental Table S8. Specificity of the respective amplification products was validated by Sanger sequencing (Macrogen, South Korea).

### Statistical analysis

Gene expression levels were analyzed as the dependent variable using a Generalized Linear Mixed Models (GLMM) with ‘colony’ as random effect, and ‘social context’ and ‘tissue’ as fixed effects. We used a log link function to all genes to approximate gene expression data to a Gaussian distribution, which was checked by analyzing the residuals’ distribution. Alternative link functions and data transformations were applied as necessary (see Supplemental Table S6). Given that the ‘social environment’ (presence/absence of the queen) might influence gene expression in opposite directions in different tissues, as seen for Dnmt’s expression, we performed Tukey’s post-hoc tests for all gene/tissue combinations. To investigate whether colony of origin influenced gene expression, ‘colony’ and ‘tissue’ were treated as fixed effects and ‘social environment’ as a random effect. Tukey’s post-hoc tests were performed to identify differences in gene expression between individuals from different colonies (Supplemental Table S7). The Pearson’s correlation coefficient between gene expression and differential methylation was calculated with a two-tailed test of significance. Statistical testing was performed in *R* (R Core Team, 2018) using the packages lme4, car and lsmeans, or in the GraphPad Prism 7 statistics package. For all analyses, a *p*-value < 0.05 was considered significant.

## Data access

All raw sequencing data generated in this study have been submitted to the NCBI Sequence Read Archive under accession number PRJNA714749.

## Acknowledgments

This work was supported by grants from São Paulo Research Foundation (FAPESP 2016/15881-0 and 2017/09269-3 to CAM; and 2017/09128-0 to KH), the Brazilian National Council for Scientific and Technological Development (CNPq 403646/20162 and 303401/2014-1 to KH) and Australian Research Council (DP180101696 to BO and A. Zayed).

