## Supplementary figures and images for "DNA methylation is not a driver of gene expression reprogramming in young honey bee workers"

### Supplementary Figure 2

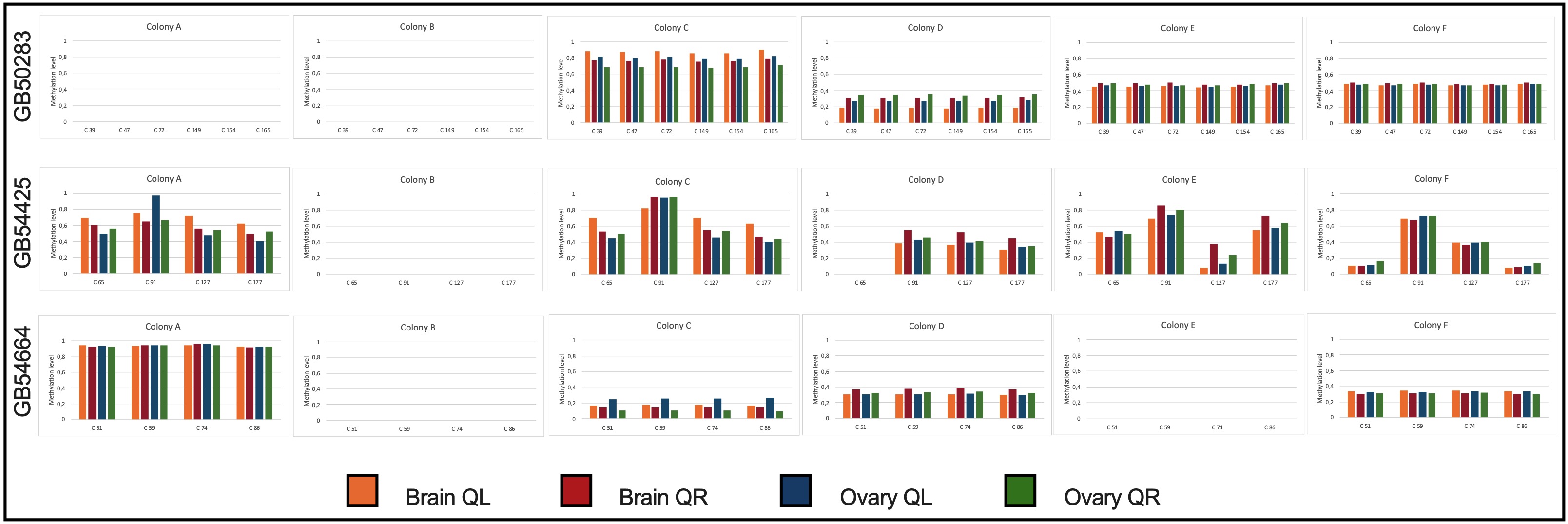

### Supplementary Figure 6

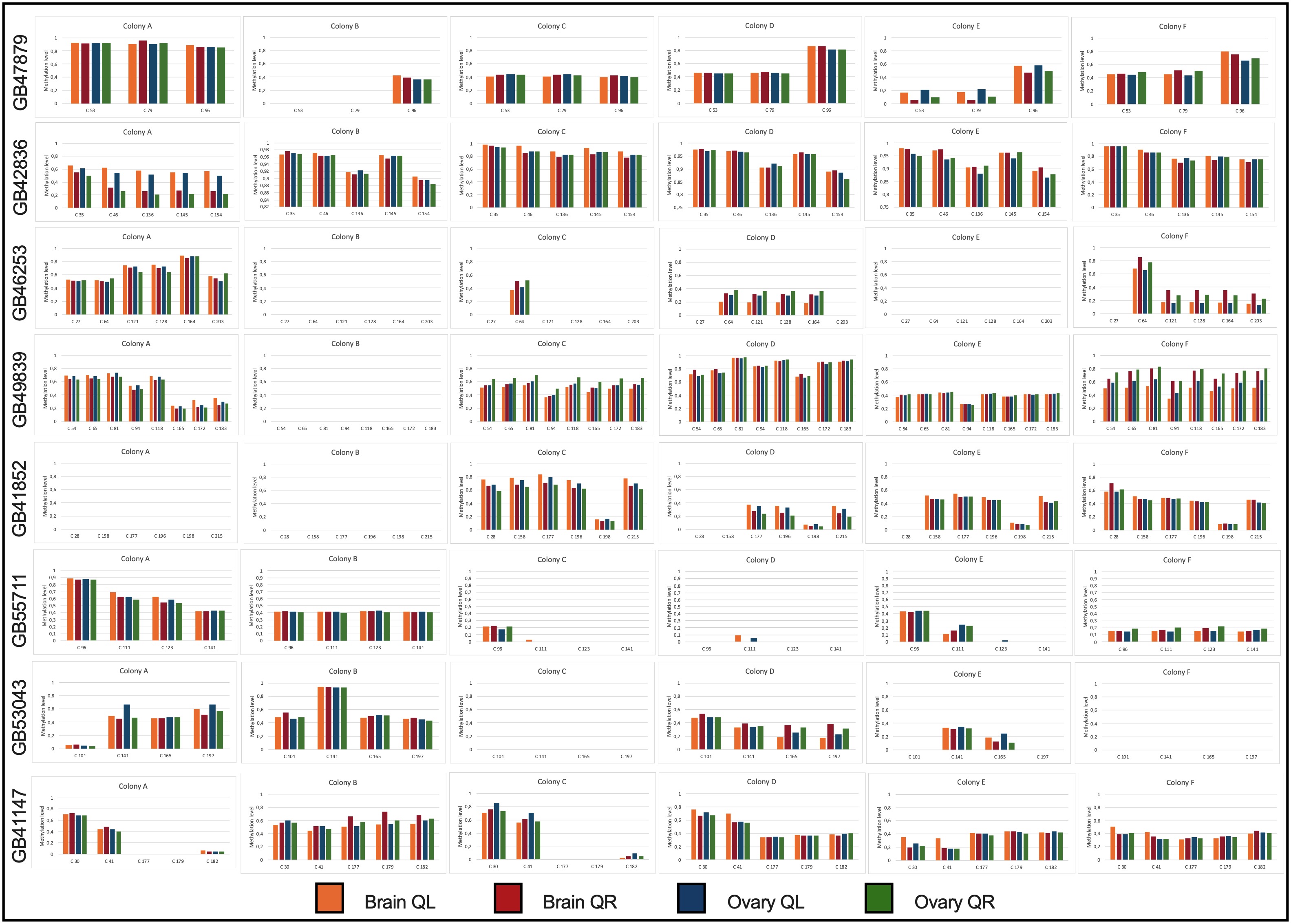

### Supplementary Figure 7

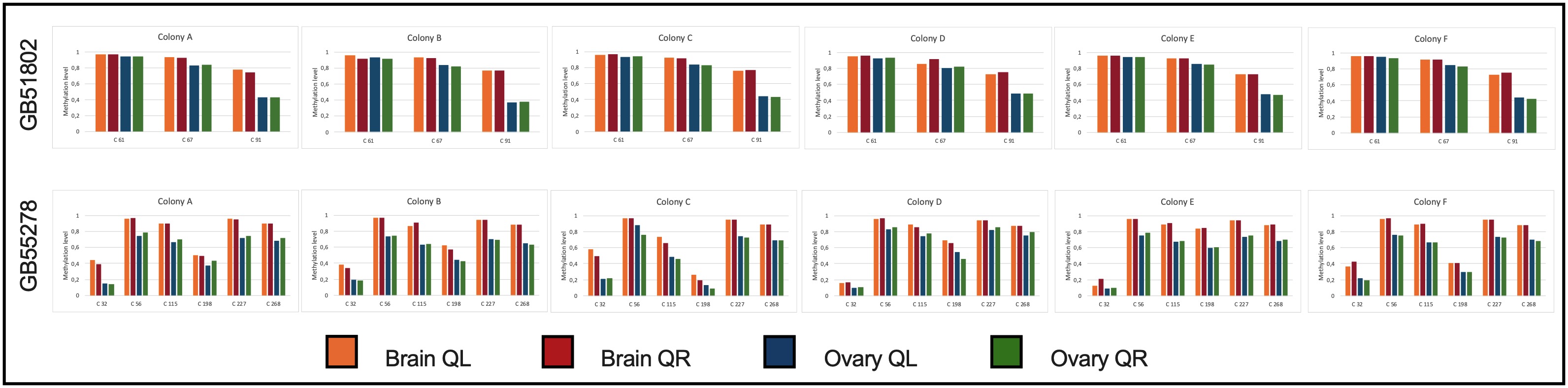
