## Supplementary Figures for "DNA methylation is not a driver of gene expression reprogramming in young honey bee workers"

Carlos Antonio Mendes Cardoso-Junior, Boris Yagound, Isobel Ronai, Emily Jane Remnant, Klaus Hartfelder, Benjamin Paul Oldroyd

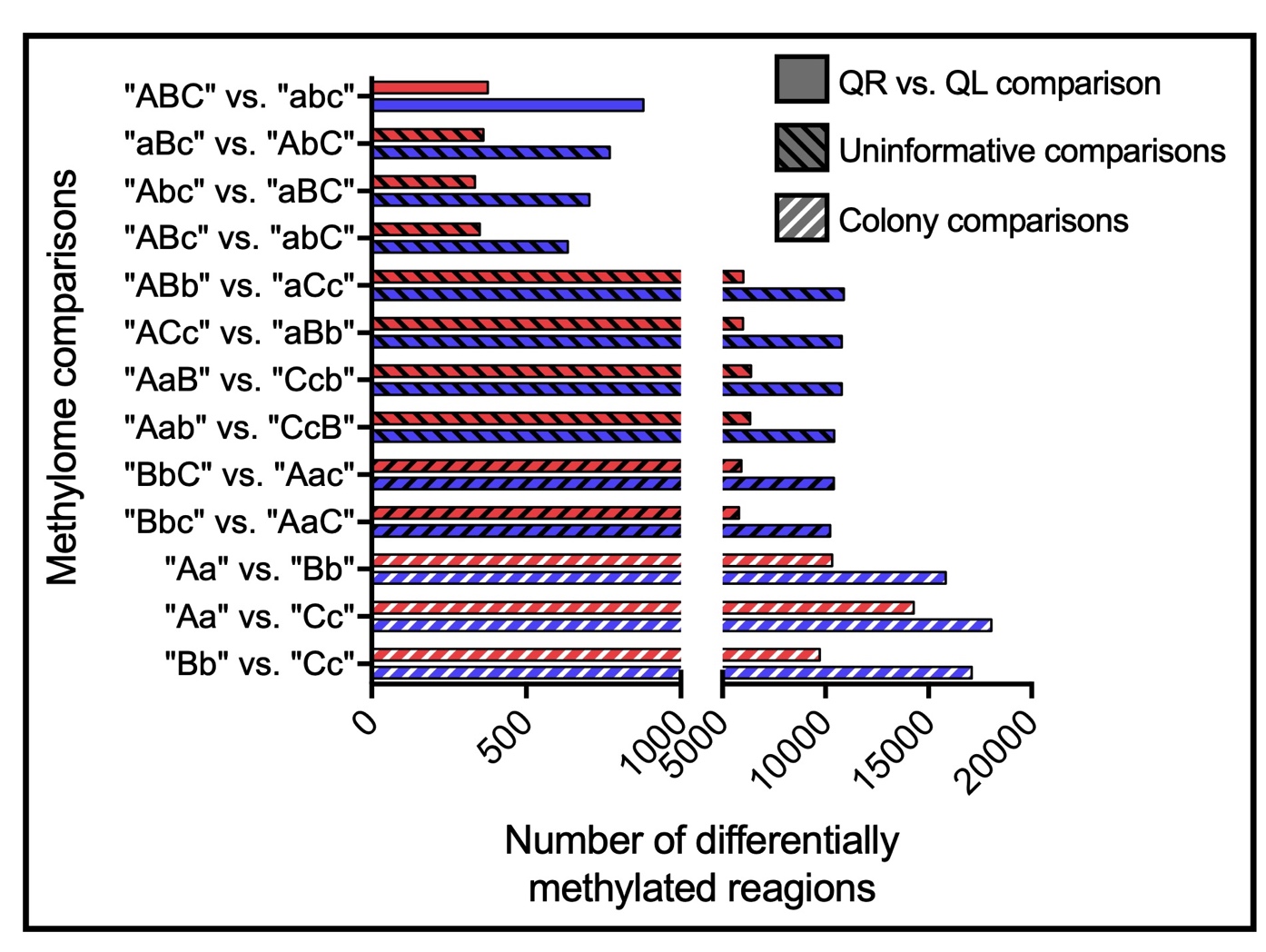

**Supplemental Fig. S1.** Number of differentially methylated regions from multiple methylome comparisons show that ‘colony’ effect is much stronger than ‘social context’ effect. A comparison of the brain (blue bars) and ovary (red bars) methylomes of workers from queenright and queenless colonies show fewer DMRs than seen in the nine uninformative combinations (black dashed bars) that lake any biological-relevant basis as they are a result of an aleatory shuffling of the samples to compare a group of three methylomes against another group of three methylomes in each tissue, not allowing repetition of samples when comparing. In contrast, a comparison of the different colonies (A *vs.* B, A *vs.* C or B *vs.* C – white dashed bars) regardless of the social environment, returned more DMRs than seen in the nine uninformative combinations.

Download separately

**Supplemental Fig. S2.** Methylation frequency validated by amplicon sequencing of CpG sites showing differential methylation between social contexts (QR *vs.* QL). DMRs were selected from WGBS data (Supplemental Tables S3 and S4). CpG sites are numbered according to the primer locations in the respective scaffold. Empty panels represent CpG methylation level detected at 0%.

Coverage is shown in Supplemental Table S5.

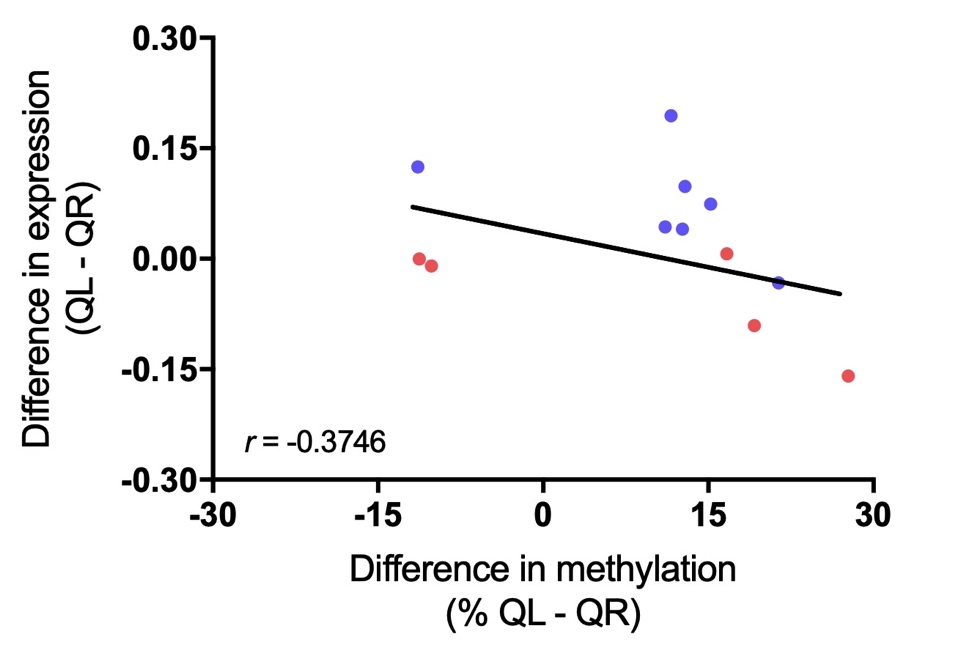

**Supplemental Fig. S3.** Correlation analysis between the differential methylation (Δ = %QL - %QR) and expression (Δ = QL - QR) for the twelve regions differentially methylated shown in Fig. 2 (Two-tailed Pearson correlation, *p* = 0.2302, N = 12). The black line represents the trend line, blue dots represent the expression of differentially methylated genes in the brains and red dots represent the expression of differentially methylated genes in the ovaries.

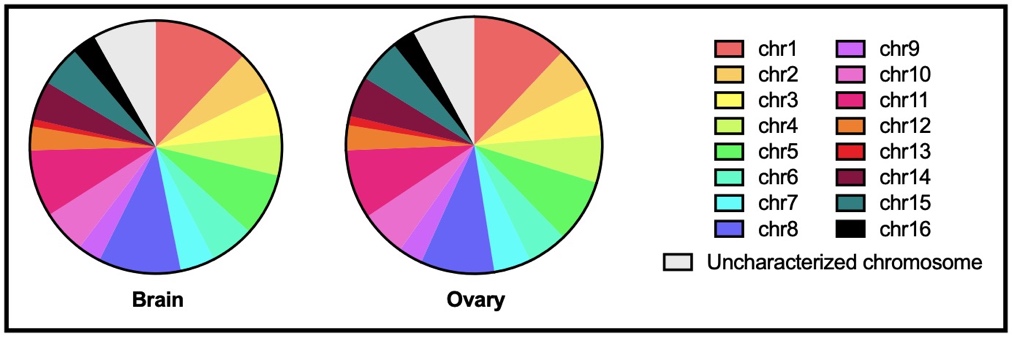

**Supplemental Fig. 4.** Genomic location of regions displaying at least 10% of differential methylation (e.g., hypervariable regions shown in Fig. 3A) comparing the methylation patterns between all three colonies analyzed by WGBS (colonies A-C).

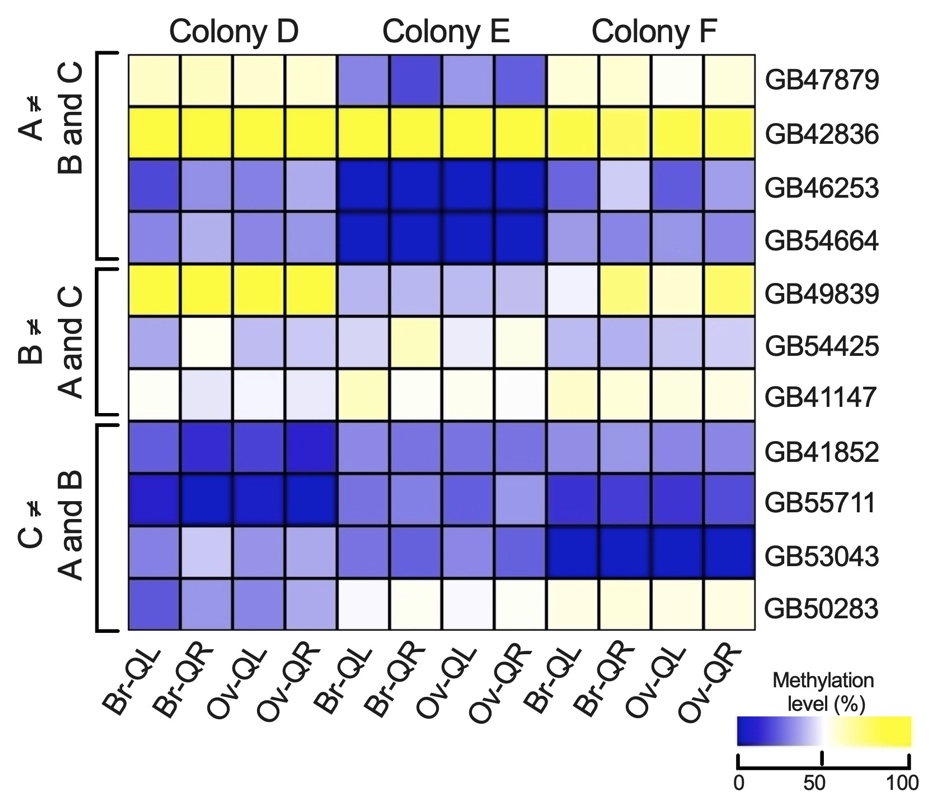

**Supplemental Fig. S5.** Methylation patterns validated by amplicon sequencing of regions displaying colony-specific methylation for colonies D-F. DMRs were selected from WGBS data based on the methylation patterns of colonies A-C. Methylation of individual CpGs can be found in Supplemental Figs. S2, S6 and coverage in Supplemental Table S5.

Download separately

**Supplemental Fig. S6.** Methylation frequency validated by amplicon sequencing of CpG sites showing differential methylation between different colonies. DMRs were selected from WGBS data (Supplemental Table S3 and S4). CpG sites are numbered according to the primer locations in the respective scaffold. DMRs from “*GB54664*”, “*GB50283*” and “*GB54425*” genes are displayed in Supplemental Fig. S2. Empty panels represent CpG methylation level detected at 0%.

Coverage is shown in Supplemental Table S5.

Download separately

**Supplemental Fig. S7.** Methylation frequency validated by amplicon sequencing of CpG sites showing differential methylation between the brain and ovary methylomes. DMRs were selected from WGBS data. CpG sites are numbered according to the primer locations in the respective scaffold. Coverage is shown in Supplemental Table S5.
